# A Magnetic Field Prevents Alcoholic Liver Disease by Reducing Oxidative Stress

**DOI:** 10.1101/2021.12.07.471564

**Authors:** Chao Song, Hanxiao Chen, Biao Yu, Lei Zhang, Junjun Wang, Chuanlin Feng, Xingxing Yang, Xiaofei Tian, Yixiang Fan, Xinmiao Ji, Hua Wang, Can Xie, Xin Zhang

## Abstract

The radical-pair recombination change will affect the generation of free radicals, which can be regulated by static magnetic fields (SMFs) in a SMF setting dependent way. It is well known that alcohol consumption leads to significantly increased free radical levels and health risks, which lacks effective treatment method besides alcohol abstinence. Here we compared different SMF settings and found that a downward SMF of ∼0.1 T with magnetic flux of ∼4.5×10^−3^ Wb could effectively alleviate alcohol-induced liver damage and lipid accumulation, and improve liver function. The inflammation, reactive oxygen species (ROS) level and oxidative stress were significantly reduced. EPR (electron paramagnetic resonance) experiments also confirmed the reduced amount of free radical by SMF treatment. Moreover, the lifespan of heavy alcohol drinking mice was also significantly changed due to the SMF effects on liver cell ROS level, DNA synthesis and liver cell regeneration. Our study shows that moderate SMFs with specific parameters have great promises to be developed into a physical method to reduce alcohol-induced liver damage in the future.

## Introduction

Alcoholic liver disease (ALD) caused by excessive drinking is one of the most common chronic liver diseases, including steatosis, steatohepatitis, fibrosis, cirrhosis and hepatocellular carcinoma. ALD with a high short-term mortality has become a severe health threat and global burden(Bhandari et al., 2020; Liang et al., 2015; Rehm et al., 2013; Witkiewitz et al., 2019; Xiao et al., 2020), especially in current COVID-19 pandemic situation(Sy-Janairo and Cua, 2020; Williams et al., 2021). A recent survey in about 7528 alcoholics reveals that 27% of them have liver steatosis, 20% have steatohepatitis and 26% have cirrhosis(Parker et al., 2019). In fact, around 2 million people die from liver disease worldwide each year and 50% of them have advanced liver cirrhosis(Szabo et al., 2019). However, although corticosteroids (prednisolone and prednisone) and pentoxifylline can be used to increase short-term survival of severe alcoholic hepatitis(Singal et al., 2018), there is no US Food and Drug Administration (FDA) approved drugs for ALD prevention or treatment besides abstinence from alcohol and subsequently nutritional supplements. Therefore, people are still actively seeking effective and safe therapies for the large number of ALD patients.

Increased free radical level and oxidative stress in liver cells play a central role in the development of alcoholic liver disease(Albano, 2006; Das and Vasudevan, 2007; Wu and Cederbaum, 2003). Magnetic field can control the movement and transfer of electrons, providing a non-invasive physical method to manipulate the unpaired electrons in free radicals, which provides a theoretical basis for cellular ROS regulation by externally applied magnetic field(Ikeya and Woodward, 2021; Timmel et al., 1998). In fact, it has been shown in multiple studies that static magnetic fields (SMFs) can affect ROS levels in many cells and animals, but the effects are variable(Wang and Zhang, 2017). Van Huizen et al. showed that the ROS levels can be affected by SMFs in a field intensity dependent way(Van Huizen et al., 2019) and Gurhan et al. confirmed that different directions had various effects on ROS change, mitochondrial calcium and cell growth in fibrosarcoma cell HT-1080(Gurhan et al., 2021). Moreover, our previous study shows that SMFs can affect ROS levels in cell type dependent manner (Wang and Zhang, 2019). Therefore, the effects of SMFs on ROS levels are closely associated with magnetic field intensity, direction and biological samples examined.

In recent years, multiple types of electromagnetic fields have been developed as non-invasive physical tool to treat cancer, depression and diabetes, including tumor treating field (TTF), transcranial magnetic stimulation (TMS) as well as static magnetic and electric fields. For example, recently, Carter et al. show that a 3 mT static magnetic field in combination of a static electric field can treat type 2 diabetes (T2D) through ROS and redox state regulation(Carter et al., 2020). Our previous study shows that a downward SMF of ∼0.1 T can reduce ROS levels in pancreatic cells and alleviate T2D(Yu et al., 2021). Van Huizen et al. show that the ROS level in Planaria regeneration can also be decreased by 200 mT SMF to alter stem cell–mediated growth(Van Huizen et al., 2019). In addition, Yu et al. found that static magnetic field combined with vesicles could promote ferroptosis-like cancer cell death(Yu et al., 2020). Therefore, modulating ROS and oxidative stress by electromagnetic field could potentially provide a non-invasive physical tool to regulate physiological and pathological processes.

In this study, we sought to examine the effects of SMFs in alcohol-induced liver damage by using different mice models and SMF settings. We found that for both shorter-term lighter drinking mice and longer-term heavier drinking mice, the alcohol-induced liver damage can both be significantly relieved by a downward SMF of ∼0.1 T with magnetic flux of ∼4.5×10^−3^ Wb. The lifespan of the heavy drinking mice was significantly prolonged by this magnetic field exposure. In contrast, a different SMF setting did not have such effects, and even be detrimental to heavy drinking mice.

## Results

### The downward SMF alleviates alcohol-induced mice liver damage

To assess the effects of SMF on alcohol-induced ALD, we used magnetic plates with different settings. The magnet plates were composed of twelve closely connected N38 Neodymium (NdFeB) magnet cubes, with either the North or South pole facing up, which provided a relatively uniform magnetic flux density at a given horizontal plane (Figure 1A). We measured the magnetic field intensity at 2 cm above the magnet plate, which corresponds to the position of the mice bodies when they stand. The average magnetic field intensity of the three groups at this horizontal level are 2.95 mT, 109.0 mT, and 111.8 mT, respectively. And the magnetic flux is ∼1.18×10^−4^ Wb, 4.36×10^−3^ Wb and 4.47×10^−3^ Wb, respectively (Figure 1A). To reduce any potential placebo effects, we used un-magnetized NdFeB as sham control. The whole mice cages were placed on the top of these magnetic plates (Figure 1B). For the mice that were exposed to the magnetized plates, the magnetic flux density on their bodies were ∼0.08-0.19 T (Figure 1B). We fed mice with the Lieber-DeCarli alcohol liquid diet(Lieber et al., 1989) containing either ethanol (EtOH-fed) or maltodextrin diet as a control (pair-fed) with unlimited access (Figure 1C). For the EtOH-fed group, we gradually increased the concentration of EtOH, from 1% (week 1), 2% (week 2) to 3% (week 3). We started the alcohol feeding and SMF exposure simultaneously from the beginning and continued for 3 weeks (SMF exposure was 12 h/d, 7 d/week) (Figure 1C).

**Figure 1.**
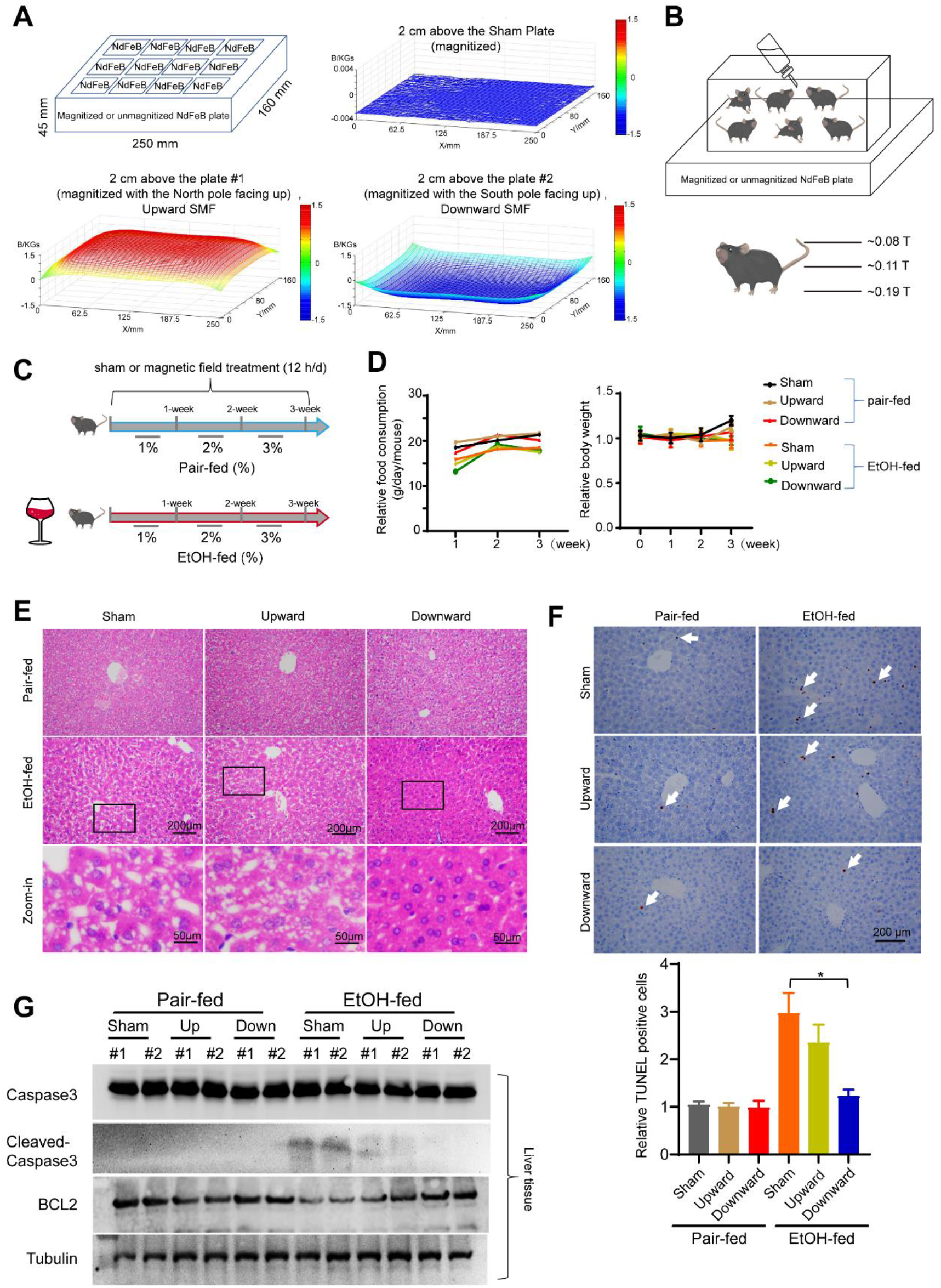
The downward static magnetic field alleviates alcohol-induced mice liver damage. **(A)** Unmagnetized or magnetized NdFeB plates were used to provide sham, upward or downward SMFs. Magnetic flux densities were scanned at 2 cm above the plates with a magnet analyzer. **(B)** The whole mice cage was place on the magnetic plate and the SMF on mice was ∼0.08-0.19 T. **(C)** The mice were fed with unlimited access of the Lieber-DeCarli alcohol liquid diet containing either ethanol (EtOH-fed) or maltodextrin diet (pair-fed) with exposure to sham, upward or downward SMFs for 3 weeks. **(D)** Food consumption and body weight were recorded every week. **(E)** Liver sections were subjected to H&E staining. **(F)** Liver section TUNEL assay and its quantification. Arrows indicate positively stained cells. n=3-5 per group. **(G)** The protein levels of cleaved-caspase 3, caspase 3, BCL2 and tubulin were detected by Western blot. All values represent means ± SEM. *, *P* < 0.05 by Student’s *t*-test or two-way analysis of variance (ANOVA) with Bonferroni correction for comparisons.

To get comprehensive information about the effect of SMFs on these mice, we monitored their body weight, food consumption and their vital signs. No significant differences in body weight or food consumption were observed among EtOH-fed groups (Figure 1D). For their vital signs, it seems that three-weeks of alcohol consumption reduced both the heart rate (from 178.2 to 160.7, *P* < 0.05) and the arterial O_2_ (from 97.6% to 93.96%, *P* < 0.05). However, it is interesting that both the SMFs can increase the mice arterial O_2_ (Figure 1-figure supplement 1).

Next, H&E staining were used to analyze the mice tissue. No significant difference was observed in spleen, lung, kidney or heart (Figure 1-figure supplement 2). However, all EtOH-fed mice had obvious liver damage, which appear as a large number of vacuoles on the liver sections. It is interesting that although both the upward and downward SMFs reduced the vacuoles, the downward SMF had a much more obvious alleviation effect (Figure 1E). Similarly, although both the upward and downward SMFs reduced the ethanol-induced liver cell apoptosis, the downward SMF had a much more significant alleviation effects as shown by the TUNEL assay (Figure 1F). Consistently, Western Blot analysis also shows that the alcohol-induced cleaved-caspase 3 was reduced by SMF treatment, especially by the downward SMF. The alcohol induced BCL2 reduction was also reversed by SMF treatment, especially by the downward SMF (Figure 1G).

### The downward SMF significantly alleviates alcoholic fatty liver

Fatty liver is a common symptom for ALD patient. To evaluate the effects of SMFs on alcohol-induced fatty liver, we used oil red O staining to stain fat deposition in liver tissue. It was obvious that the downward SMF significantly reduced lipid accumulation in liver of the EtOH-fed mice, while the upward SMF had no obvious effects (Figure 2A). Moreover, the alcohol-induced serum alanine aminotransferase (ALT), aspartate aminotransferase (AST), Triglycerides (TG) elevation, as well as high density lipoprotein cholesterol (HDL-c) reduction, were all significantly alleviated by the downward SMF treatment but not the upward SMF (Figure 2B, C). These observations suggest that the downward SMF can improve lipid metabolism and reduce fatty liver in EtOH-fed mice.

**Figure 2.**
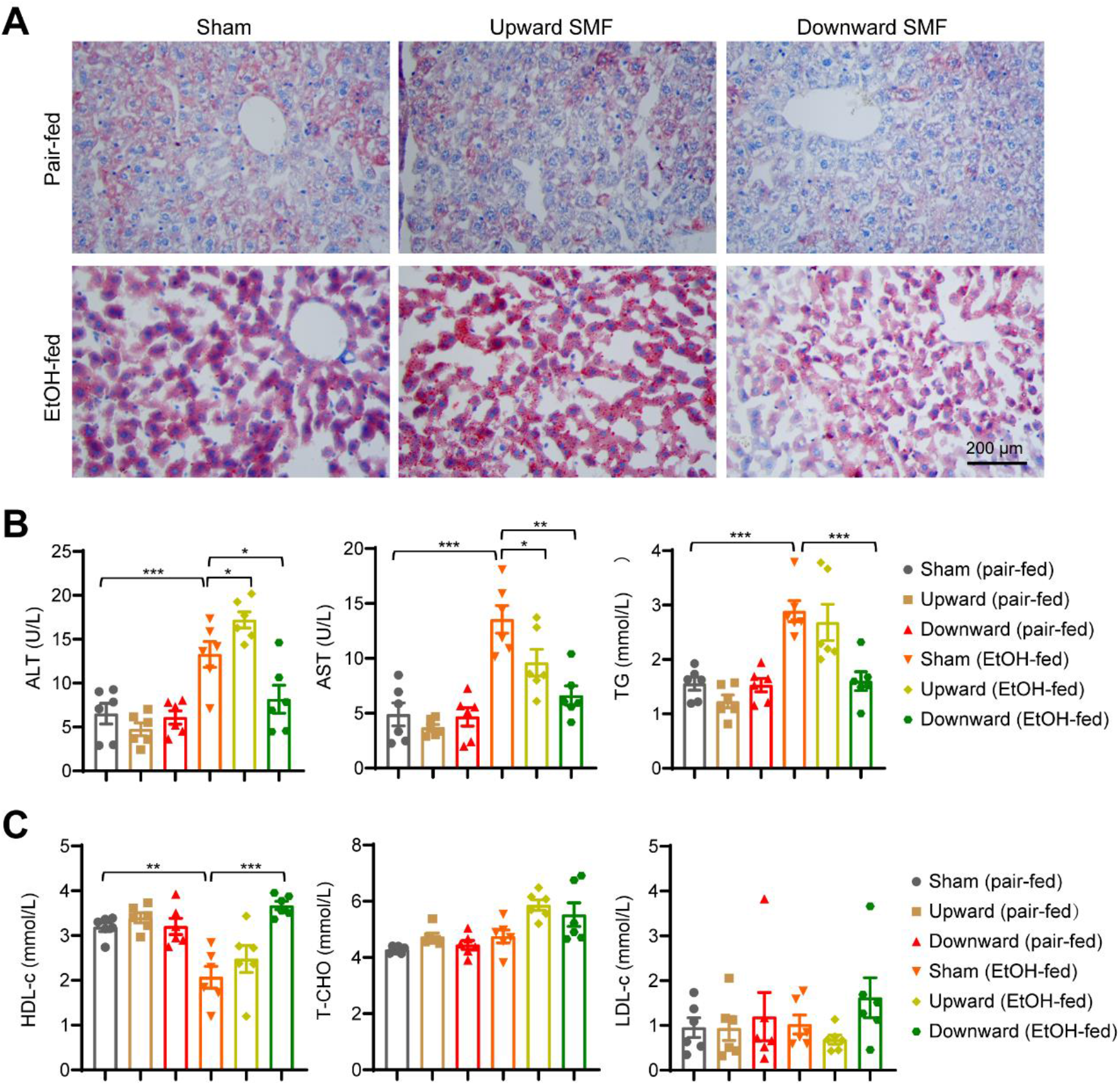
The downward static magnetic field significantly alleviates alcoholic fatty liver. **(A)** Paraffin-embedded liver sections were used to assess lipid accumulation by oil red O staining. Scale bar: 200 µm. **(B)** Serum ALT, AST and triglyceride (TG) were measured. **(C)** Serum HDL-c, LDL-c and T-CHO were measured. Values represent means ± SEM, n=6 per group. *, *P* < 0.05; **, *P* < 0.01; ***, *P* < 0.001 by Student’s *t*-test.

### The downward SMF alleviates alcohol-induced inflammation and oxidative stress in liver

To examine liver inflammation, one of the main characters of ALD, we first used F4/80 immunohistochemistry staining to identify Kupffer cells, the liver resident macrophages that play important role in liver inflammatory responses. Consistent with previous study(Wang et al., 2020), the Kupffer cells were increased in EtOH-fed mice (Figure 3A, B). Moreover, the downward SMF significantly reduced the Kupffer cell numbers in EtOH-fed mice (Figure 3A, B). Furthermore, we used myeloperoxidase (MPO) antibody to identify neutrophils that were recruited to the liver in response to ethanol consumption (Figure 3C, D). It was obvious that the downward SMF could also reduce the MPO positive neutrophils in EtOH-fed mice (Figure 3C, D). We also examined the mRNA levels of pro-inflammatory cytokines in mice liver, including IL-1β, IL-6, MCP and TNFα-1. Significant cytokine mRNA level reductions were observed in EtOH-fed mice exposed to the downward SMF compared to sham group (Figure 3E). These results show that the downward SMF can alleviate alcohol-induced liver inflammation.

**Figure 3.**
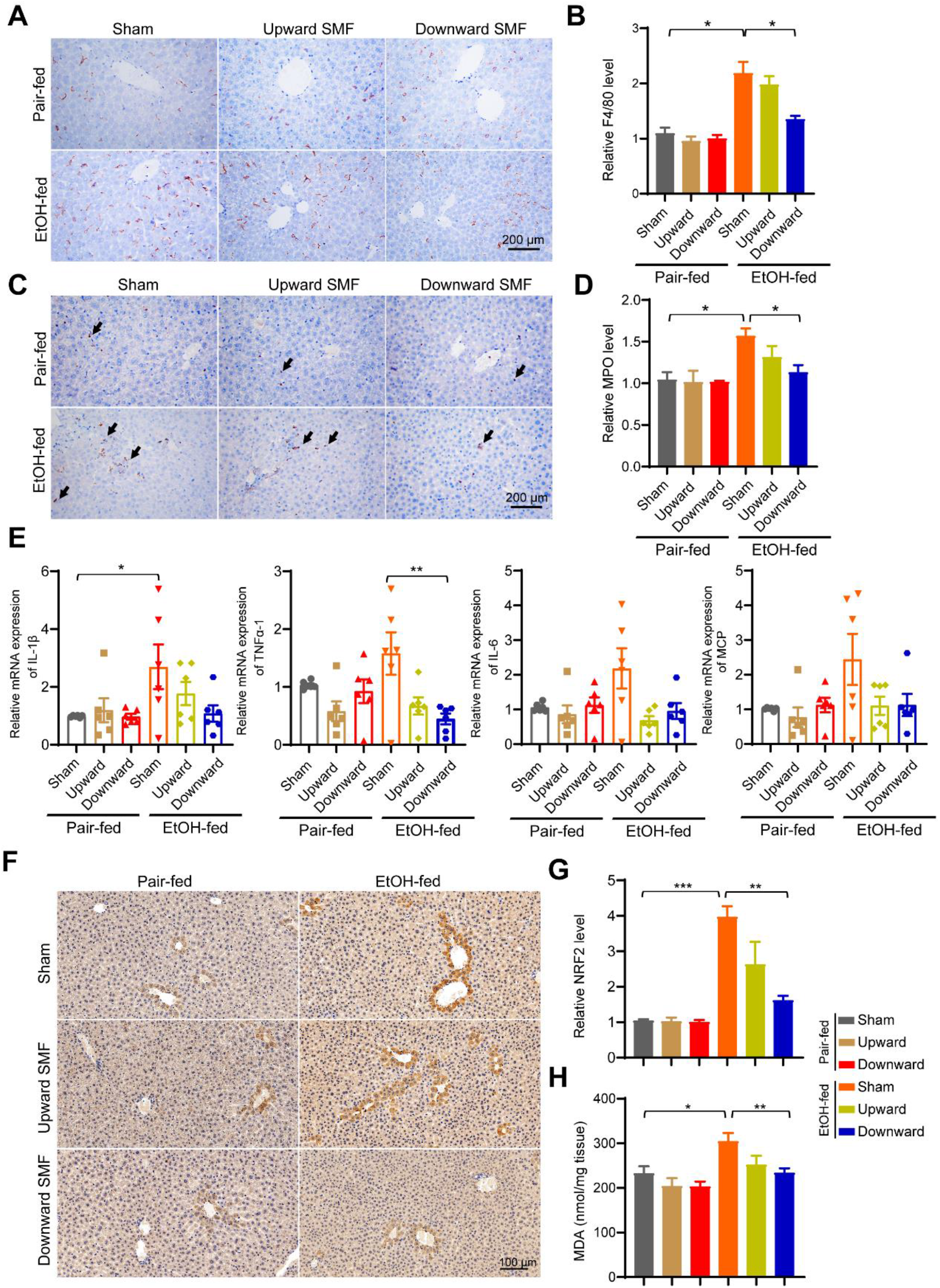
The downward SMF alleviates alcohol-induced inflammation and oxidative stress in liver. **(A, B)** Liver section immunohistochemical analysis of F4/80 positive cells and their quantification. **(C, D)** Liver section immunohistochemical analysis of MPO positive cells and their quantification. **(E)** The mRNA expressions of pro-inflammation factors were detected by real-time fluorescence quantitative PCR, including IL-1β, IL-6, MCP and TNFα-1. **(F, G)** Liver section immunohistochemical analysis of NRF2 and their quantification. **(H)** Malondialdehyde (MDA) in liver tissues were detected by the lipid peroxidation MDA assay kit. All values were expressed as means ± SEM. n=3-5 per group. *, *P* < 0.05 by Student’s *t*-test.

Since oxidative stress plays a central role in the development of alcoholic liver disease and magnetic field can affect the ROS levels in multiple studies, we next examined the oxidative stress state by using two oxidative stress indicators, NRF2 and malondialdehyde (MDA). As expected, both NRF2 and MDA levels were increased in EtOH-fed mice (Figure 3F-H). Moreover, the upregulated NRF2 and MDA in EtOH-fed mice were obviously reduced by the downward SMF treatment (Figure 3F-H). These results indicate that EtOH-induced oxidative stress can be reduced by the downward SMF treatment.

### The downward SMF protects hepatocyte through suppressing oxidative stress

To investigate the effects of SMFs on the oxidative stress in hepatocyte, we exposed HL7702 cells to NdFeB magnets that can be placed in the regular cell incubator and their magnetic field flux density is similar to the mice SMF exposure system (Figure 4A). For cells on the magnet, the SMFs are in the range of 0.1∼0.4 T.

**Figure 4.**
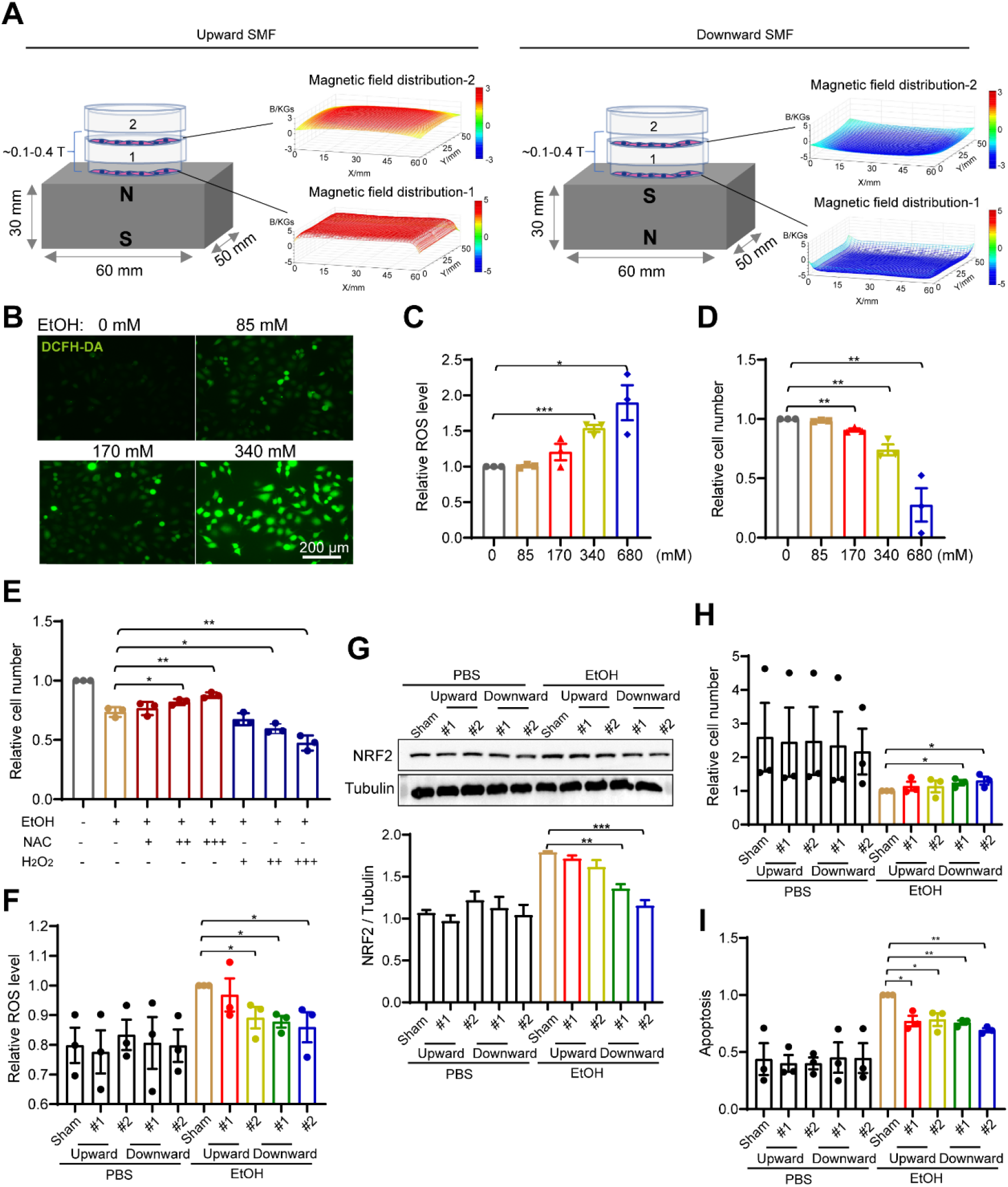
The downward static magnetic field protects hepatocyte through suppressing oxidative stress. **(A)** Experimental setup and magnetic field distribution for cells exposed to upward and downward SMFs. **(B)** Representative fluorescent images of cellular ROS levels in 24 h EtOH-treated hepatocyte HL7702 cells using a DCFH-DA probe. **(C)** Flow cytometry measurement of ROS levels in 24 h EtOH-treated hepatocyte HL7702 cells using a DCFH-DA probe. **(D)** Cell number of HL7702 cells after 24 h EtOH-treatment. **(E)** HL7702 cells (4 ×10^4^ cells/mL) exposed to 340 mM ethanol were treated with NAC (“+”, 2.8 mM; “++”, 5.6 mM; “+++”, 11.2 mM) or H_2_O_2_ (“+”, 20 µM; “++”, 30 µM; “+++”, 40 µM) for 24 h before their cell numbers were counted by flow cytometer. **(F)** Flow cytometry measurement of cellular ROS of HL7702 cells exposed to 340 mM ethanol treated with sham, upward or downward SMFs for 24 h. **(G)** Western blot analysis and quantifications of NRF2 in HL7702 cells exposed to ethanol and SMFs. **(H)** Cell number quantification by flow cytometer of HL7702 cells (4 ×10^4^ cells/mL) exposed to 340 mM ethanol and sham, upward or downward SMFs for 24 h. **(I)** Apoptosis analysis by flow cytometer using the annexin V-FITC / PI stain of HL7702 cells exposed to 340 mM ethanol and sham, upward or downward SMFs for 24 h. All data were normalized, and values were expressed as means ± SEM, n=3-5 per group. *, *P* < 0.05; **, *P* < 0.01; ***, *P* < 0.001 by Student’s *t*-test or one-way analysis of variance (ANOVA) with Bonferroni correction for comparisons.

We first confirmed that the ROS levels of the hepatocyte HL7702 cells were dose-dependently increased by EtOH treatment (*P* < 0.05, *F*= 9.76) (Figure 4B, C). Moreover, the cell numbers were dose-dependently decreased by EtOH treatment ((*P* < 0.05, *F*= 20.63), confirming the toxicity of EtOH treatment (Figure 4D). To examine the role of cellular ROS on EtOH-induced cell number reduction, we used both the ROS scavenger, N-Acetyl-L-cysteine (NAC), and H_2_O_2_ to decrease or increase the ROS levels, respectively (Figure 4E). Our results show that NAC could reduce the toxicity of EtOH while H_2_O_2_ further increased the HL7702 hepatocyte cell number reduction effects of alcohol (Figure 4E).

In order to analyze the effect of SMF on EtOH-induced cellular ROS elevation, we treated HL7702 hepatocytes that were exposed to SMFs with 340 mM ethanol for 24 h. The downward SMF significantly reduced EtOH-induced cellular ROS while the upward SMFs were less effective (Figure 4F). Since NRF2 is an oxidative stress marker, which may active antioxidant enzyme to reduce intracellular ROS caused by alcohol-induced oxidative stress(Wang et al., 2014), we next examined the effect of SMF on cellular NRF2 level. Consistent with the liver tissue results, Western blot results and their quantifications confirmed that EtOH-increased cellular NRF2 protein level was significantly decreased by the SMFs, especially by the downward SMF (Figure 4G). Furthermore, the EtOH-induced hepatocyte cell number reduction (Figure 4H) and apoptosis (Figure 4I) can both be inhibited by SMFs, especially the downward SMF. These data suggested that the downward SMF could reduce cytotoxicity by reducing EtOH-induced cellular ROS and oxidative stress.

### The downward SMF effectively reverses the alcohol- and H_2_O_2_-induced lipid accumulation

To examine the effect of SMF on liver cell lipid accumulation, we used oil red O to stain HL7702 cells. Consistent with the mice liver tissue results, the oil red O staining shows that alcohol could dose-dependently increase cellular lipid accumulation (*P* < *0*.*05, F*= 10.59) (Figure 5A, B). Moreover, the lipid accumulation could be significantly reversed by NAC, or aggravated by H_2_O_2_, which demonstrated the correlation between ROS levels and lipid accumulation (Figure 5C, D). In addition, the downward SMF not only obviously reduced lipid accumulation in cells that were treated by alcohol, but also by H_2_O_2_ (Figure 5E, F). This suggests that the downward SMF could reduce ROS levels and prevent alcohol or other oxidative stress-induced fat deposition in hepatocyte.

**Figure 5.**
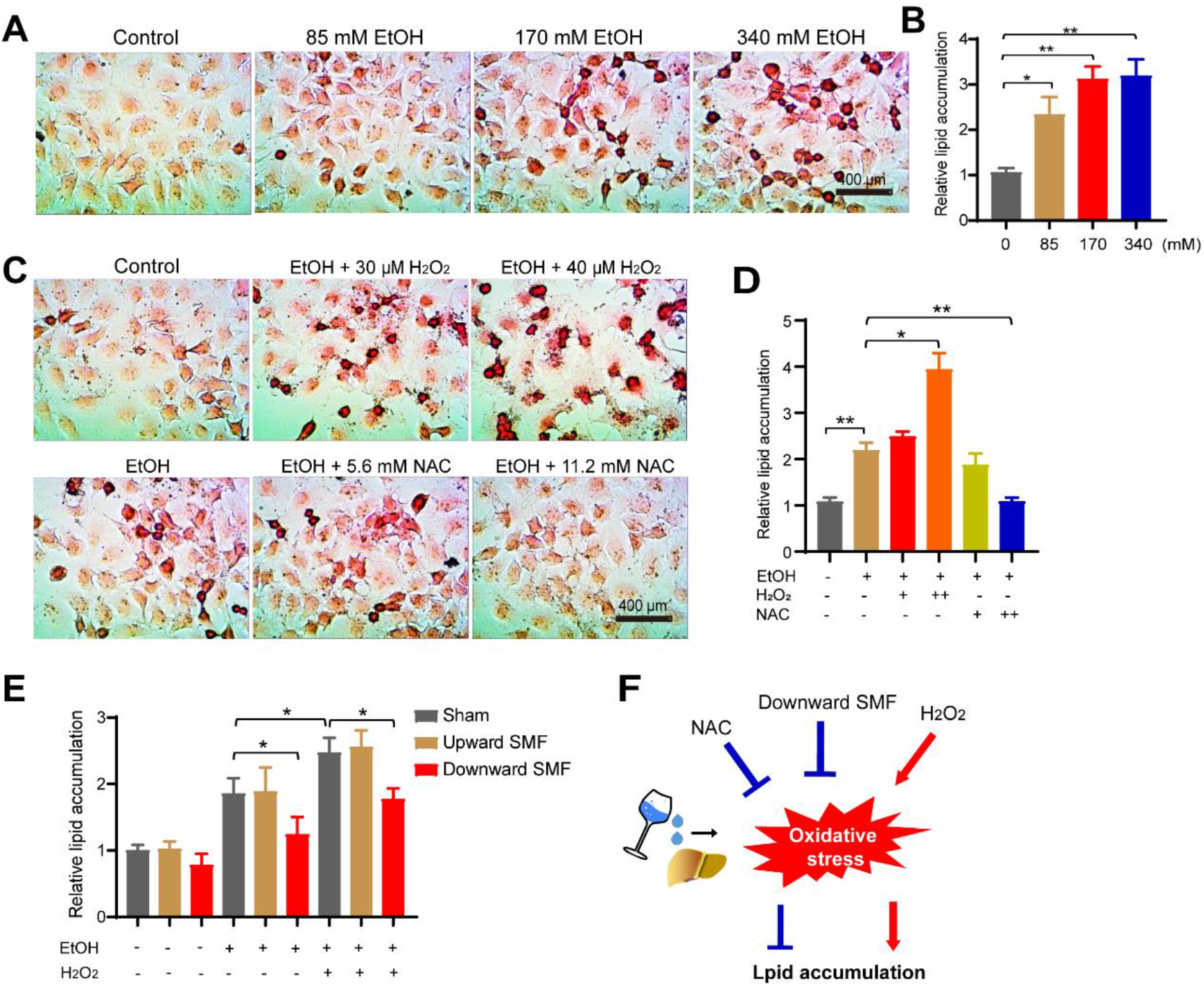
The downward static magnetic field effectively reverses the alcohol-and H_2_O_2_-induced lipid accumulation. **(A, B)** Representative images of oil red O staining and quantifications of HL7702 cells (4 ×10^4^ cells/mL) treated with different concentrations of ethanol for 24 h. **(C, D)** Representative images of oil red O staining and quantifications of HL7702 cells (4 ×10^4^ cells/mL) treated with 340 mM ethanol and different concentrations of NAC or H_2_O_2_ for 24 h. **(E)** Quantifications of lipid accumulation in HL7702 cells (4 ×10^4^ cells/mL) treated with 340 mM ethanol and upward or downward SMFs for 24 h. **(F)** Mechanism illustration of the effect of downward SMF on hepatocyte oxidative stress and lipid accumulation. Values were expressed as means ± SEM, n=3-5 per group. *, *P* < 0.05; **, *P* < 0.01 by Student’s *t*-test or one-way analysis of variance (ANOVA) with Bonferroni correction for comparisons.

### The downward SMF obviously reduces free radicals in EPR (electron paramagnetic resonance) experiment

To further confirm the effect of SMF on oxidative stress and ROS, we detected the changes of free radicals in H_2_O_2_ solution when exposed to different SMF conditions (Figure 6A, B). 0.2 M H_2_O_2_ solution was placed at different positions on the surface of the magnet for 24 h, which have different magnetic field distributions. The magnets are the same as in Figure 4A. It is obvious that the SMF at position #a is uniform while the SMF at position #b is gradient. The commonly used spin trap, 5,5-dimethyl-1-pyrroline N-oxide (DMPO), were employed for EPR free radical detection. The results showed that the free radical levels were decreased by the SMF treatment (Figure 6C, E). In particular, we identified an obvious reduction in the hydroxyl radical (OH·) level after SMF treatment, especially the downward SMFs(Figure 6D, F). This is consistent with the more efficient ROS level reduction by downward SMFs in cellular studies (Figure 4F).

**Figure 6.**
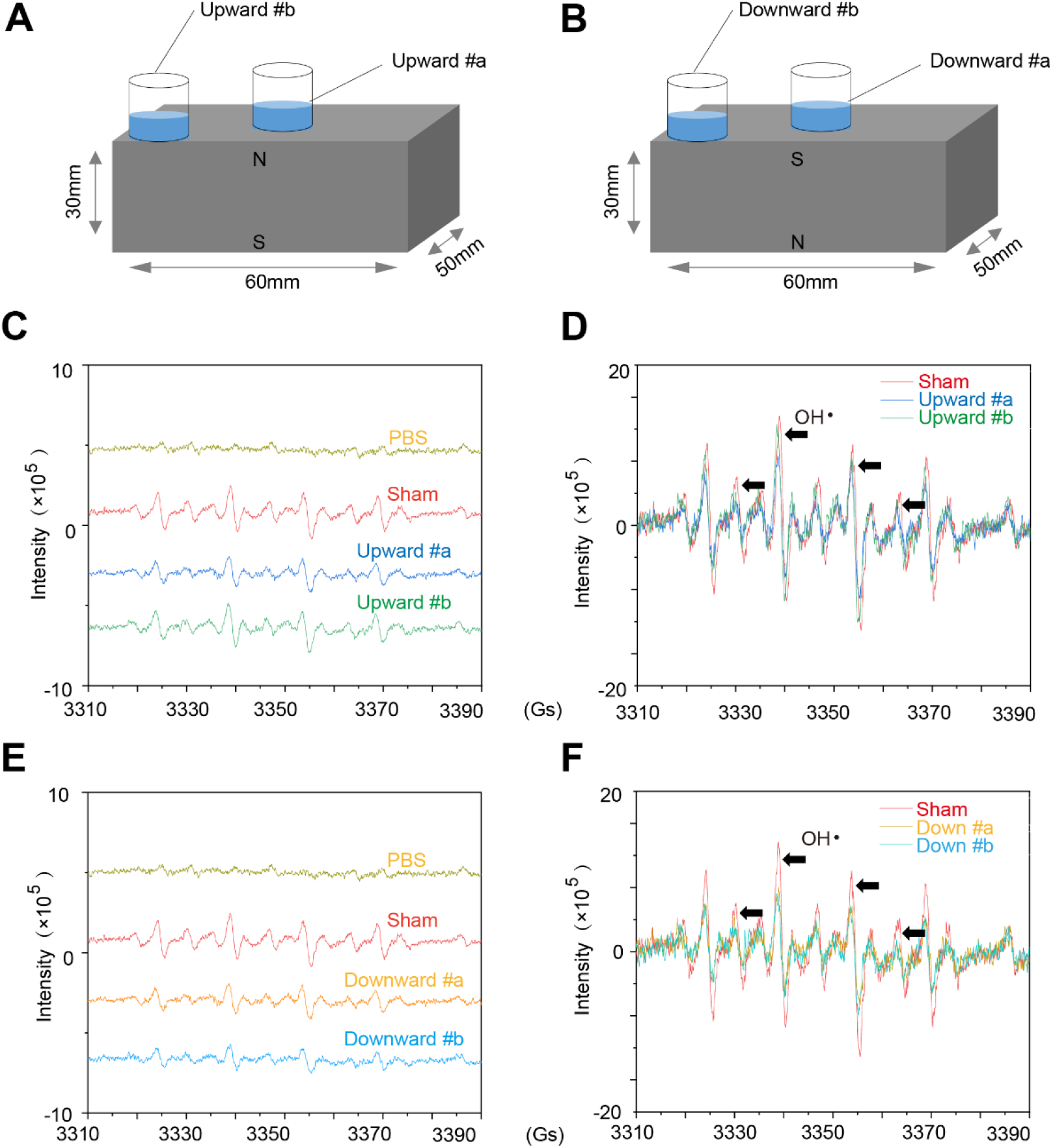
The SMFs obviously reduced free radicals in EPR experiment, especially the downward SMFs and homogeneous SMFs. **(A, B)** 0.2 M H_2_O_2_ solution were placed at different positions on the surface of two magnets. The gradient at position #b is higher than position #a. **(C, D)** EPR experiments were used to detect and analyze the level of free radicals in H_2_O_2_ solution exposed to the upward SMFs. **(E, F)** EPR experiments were used to detect and analyze the level of free radicals in H_2_O_2_ solution exposed to the downward SMFs. The black arrows indicate the spectrum of hydroxyl radical (OH·).

### The downward SMF improves the survival rate of heavy drinking mice

It is well known that excessive amount of alcohol consumption, heavy drinking or binge drinking, can cause detrimental effects and lead to high death rate(Labry et al., 1992; Esser et al., 2020; Marmot et al., 1981; Shaper et al., 1988). Our experiments above clearly show that moderate SMFs, especially the downward SMF, can reduce the oxidative stress and have obvious beneficial effects on alcohol-induced mice liver lipid accumulation and toxicity. Here, we wanted to address whether SMFs can reduce the mortality of heavy drinking mice by using an acute EtOH mice model (Wang et al., 2020) (Figure 7A). Thirty-six mice were randomly divided into six groups with either EtOH-fed or pair-fed free access Lieber DeCarli diet, with sham, upward or downward SMF treatment. The lifespan of the EtOH-fed mice in the sham group is 31±2.88 days while the pair-fed mice lived normally until the end of the experiment (Figure 7B), which confirms the detrimental effects of excessive alcohol consumption.

**Figure 7.**
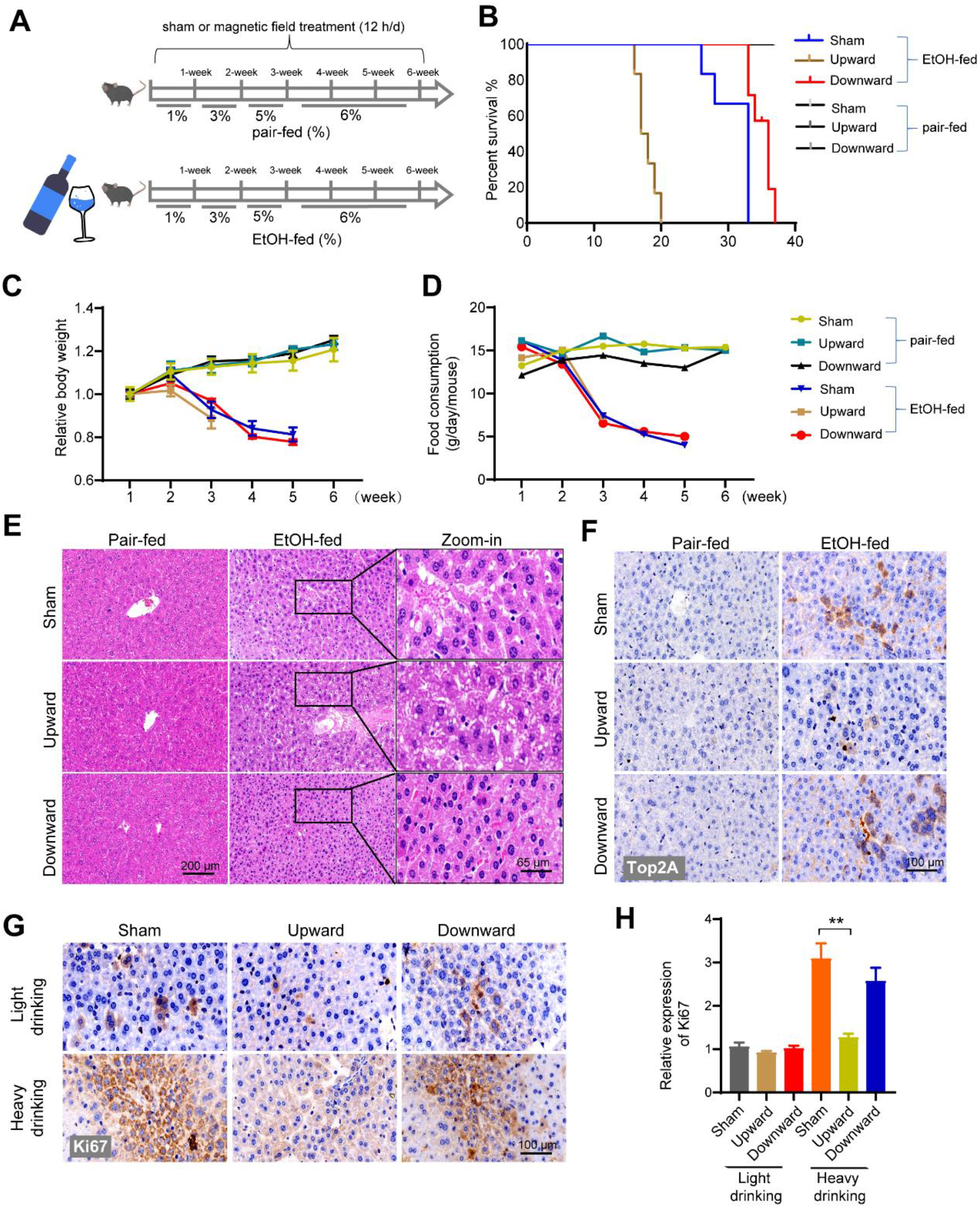
The downward static magnetic field improves the survival rate of mice exposed to alcohol. **(A)** The mice were exposed to a long-term EtOH-fed diet or pair-fed diet. **(B)** The survival rate of mice exposed to long-term diets. **(C, D)** Food consumption and body weight were recorded every week in sham, upward and downward SMFs groups. **(E)** After EtOH-fed mice died, paraffin-embedded liver sections were stained with hematoxylin-eosin (H&E). Scale bar: 100 µm and 65 µm. **(F)** The expression of Top2A were detected in liver sections. Scale bar: 100 µm. **(G, H)** Liver sections were subjected to immunohistochemistry analysis by using antibody Ki67. Quantified analysis of images by ImageJ software. Representative images are shown. **, *P* < 0.01 by Student’s *t*-test or two-way analysis of variance (ANOVA) with Bonferroni correction for comparisons.

Consistent with our previous results showing the beneficial effects of downward SMF on light drinking mice, the lifespan of excessive EtOH-fed mice (binge alcohol drinking) was prolonged from 31±2.88 to 34.83±1.57 days (*P*< 0.05) (Figure 7B). But unexpectedly, the upward direction SMF exposure generated obvious detrimental effects on these alcohol-intoxicated mice (Figure 7B), which is different from the subtle beneficial effects of the upward direction SMF exposure on light drinking mice. The lifespan is sharply reduced to 17.83+1.34 days, which is 42.5% shorter than the sham control, after upward SMF exposure (Figure 7B). All EtOH-fed mice in the upward SMF group died before day 20^th^. SMFs had no significant effects on food consumption or body weight (Figure 7C, D), which excluded the possibility that SMF-induced effects were due to increased EtOH intake.

To investigate the death reason of these heavy drinking mice, we performed H&E staining for the heart, liver, spleen, lung, and kidney of the remaining mice on day 37 (Figure 7-figure supplement 5). Consistent with previous results in light drinking mice, the serious pathological damages induced by EtOH were obviously alleviated by the downward SMF treatment (Figure 7E). Interestingly, the immunohistochemistry results of topoisomerase 2A (Top2A) (Figure 7F) and Ki67 (Figure 7G) in the liver sections show that the heavy drinking mice have significantly increased Top2A and Ki67 levels. This indicates that there were active cell proliferation and liver regeneration after alcohol-induced liver cell damage. Moreover importantly, both Top2A and Ki67 were significantly downregulated by the upward SMF (Figure 7F, G), showing that the liver cell proliferation and liver regeneration were inhibited by the upward SMF. This is consistent with our previous findings that the upward SMF can inhibit DNA synthesis by Lorentz forces exerted on the negatively charged DNA(Yang et al., 2020). We also examined the liver sections of acute heavy drinking and chronic light drinking mice for Ki67 immunohistochemistry to evaluate the liver regeneration (Figure 7-figure supplement 6). It is obvious that the Ki67 staining level in chronic light drinking mice is much lower than the acute heavy drinking mice. Consequently, the Ki67 level was not significantly affected by the upward SMF in light drinking mice, but an obvious reduction was observed in heavy drinking mice (Figure 7G, H), indicating liver regeneration was inhibited by the upward SMF, but not the downward SMF (Figure 8). This is consistent with our previous findings that the upward and downward SMFs have differential effects on the DNA supercoil and movement, which resulted in differential effects on DNA synthesis(Yang et al., 2020). Consistently, our results show that the DNA synthesis in hepatocytes was inhibited by the upward SMF (Figure 8-figure supplement 7).

**Figure 8.**
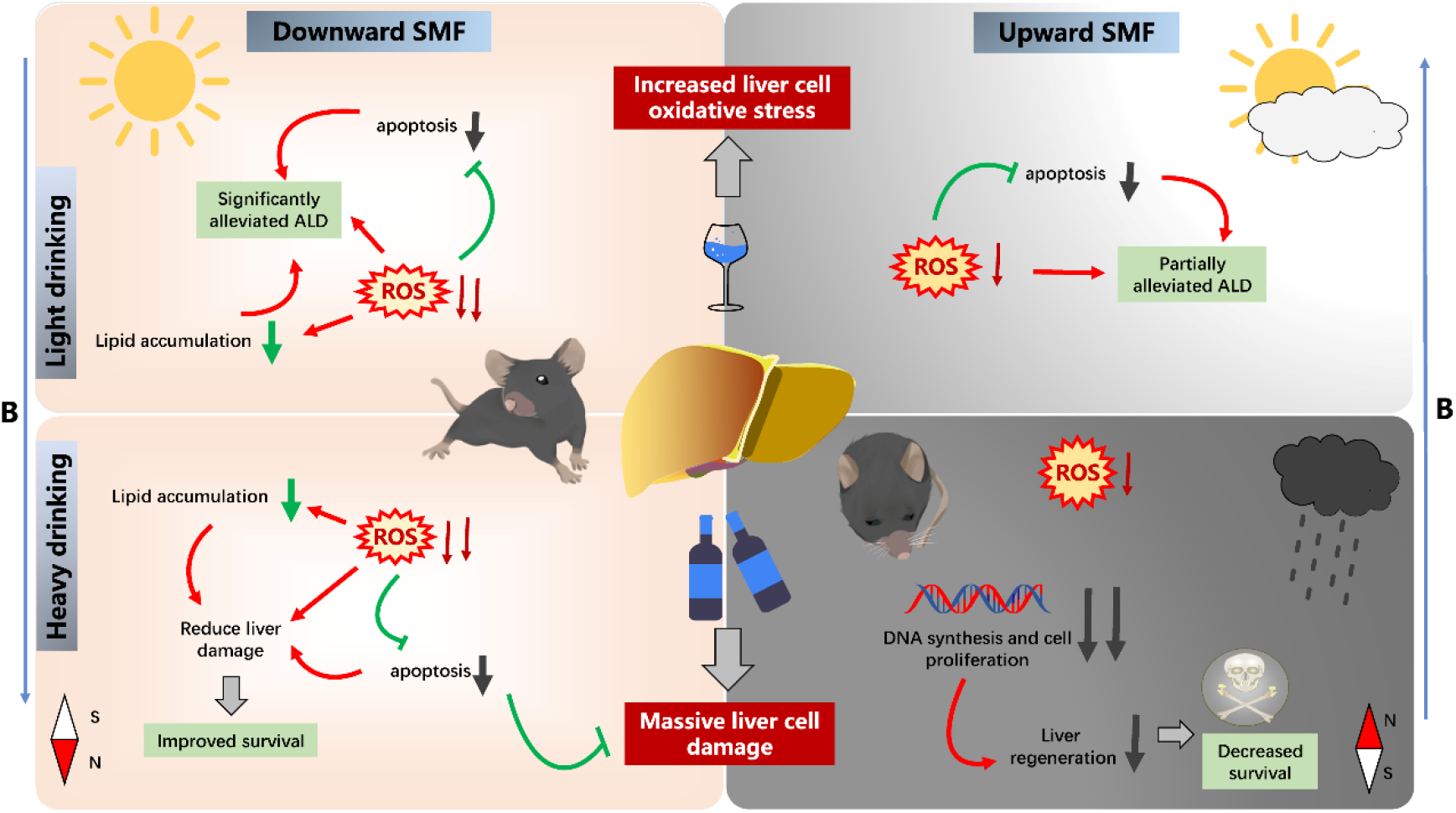
Mechanism of the upward and downward SMF treatment on light drinking vs. heavy drinking mice.

## Discussion

Although alcoholic metabolism could cause multiple organs injury(Cederbaum et al., 2012; Osna and Kharbanda, 2016), liver is still the major target for the alcohol-induced detrimental effects since the alcoholic substance is mainly metabolized by hepatocytes. However, there is no effective treatment for ALD(Liang et al., 2015), which has become a global burden on public health and economics. We found that although moderate SMFs of both upward and downward directions could reduce the liver oxidative stress and inflammation to alleviate ALD, their overall effects on the alcohol-consuming mice are different (Figure 8). The exact effects depend on the alcohol consuming amount and magnetic field direction, and likely through the radical pair mechanism.

First of all, the livers are in very different conditions in light drinking vs. heavy drinking mice. Alcohol induced cellular ROS elevation and cell number reduction in a concentration dependent way. From our cellular analysis, EtOH of 85 mM did not have significant effect on cellular ROS or cell number. However, for EtOH of 340 mM and above, the ROS elevation and cell number reduction was dramatic. In mice experiments, it was also evident that the liver of the heavy drinking mice, but not the light drinking mice, underwent obvious liver cell proliferation. This is consistent with the previous finding that the expression of Ki67 in liver was upregulated by alcohol(Alarcon-Sanchez et al., 2020).This is because the liver needs to go through liver regeneration for recovery after heavy alcohol consumption.

Secondly, the upward SMFs have more significantly inhibition effects on DNA synthesis and cell proliferation. We have previously shown that the upward SMFs could affect the DNA rotation and supercoil tightness by aligning DNA chain and generating Lorentz forces on negatively charged DNA, which inhibited DNA synthesis, but not downward SMF(Yang et al., 2020; Yang et al., 2021). This effect will lead to inhibited cell proliferation, tumor growth inhibition, as well as liver cell regeneration. Our results showed that the upward SMF decreased the expression of Ki67 and Top2A in EtOH-fed mice. We speculated that the hepatocytes regeneration was inhibited by the upward SMF in EtOH-fed mice. Therefore, the EtOH-fed mice had a low survival rate when exposed to the upward SMF.

Thirdly, although the moderate SMFs of both upward and downward directions can reduce the liver oxidative stress and inflammation, the downward SMF had much more significant effects. The oxidative metabolism of alcohol generates an obvious increase of ROS(Sha’fie et al., 2020), and ROS-induced oxidative stress is a major factor that causes liver injury(Mansouri et al., 2018). It is known that change of the radical-pair recombination will affect the generation of free radicals(Barnes and Greenebaum, 2018; Grissom et al., 1995), which can be affected by SMFs in a field intensity dependent way(Van Huizen et al., 2019). However, the exact effects of SMFs on ROS levels in cells are very complicated. For example, a given moderate SMF could affect the ROS levels in MCF7, RPE1, C6, U251, HepG2, GIST-T1, EJ1 cells, but not CHO, 293T, PC12 cells(Wang and Zhang, 2019). In fact, there are many studies that have found the influence of magnetic fields on ROS(Wang and Zhang, 2017), but people still do not have an unambiguous conclusion about the exact consequences on ROS level or the mechanisms that caused the differential effects. The ROS reduction of the upward SMFs is not as significant as the downward SMF, which is probably due to the tightened supercoil of the DNA caused by upward SMF, which induces extra stress to the cells. Furthermore, the *in vitro* study of H_2_O_2_ solution showed that the free radical level was significantly reduced by moderate SMFs, especially for the downward SMF. Interestingly, these reductions of free radicals seem to be related to the magnetic field parameters. For example, in Figure 6, both downward SMF positions #a and #b could effectively reduce the free radical levels, while the upward gradient SMF did not have obvious effect.

Fourthly, the alleviated oxidative stress and inflammation, as well as reduced lipid levels in serum and improved ALD in our study are likely to be the consequences of reducing ROS. Alcohol and its metabolites have severe hepatotoxicity, cause oxidative stress and inflammation, which significantly increased serum ALT, AST and TG in mice. It is known that the oxidative stress is mediated by cellular ROS in alcohol associated liver disease(Mansouri et al., 2018). Our results show that the downward SMF can reduce lipid accumulation and increase cell number by suppressing oxidative stress in liver, which improves ALD.

Lastly, the SMF direction dependent effects exist, but the mechanism is still elusive. We found that the downward SMF had significantly beneficial effects, including reduced liver cell apoptosis and lipid accumulation, serum ALT, AST and TG, as well as increased life span. However, the upward SMFs did not affect the light drinking mice, but surprisingly shortened the lifespan of heavy drinking mice, which is likely due to the impaired liver regeneration of the heavy drinking mice. This shares the similar mechanism with the tumor cell proliferation inhibition effects by SMFs. Moreover, we have also previously reported that the blood sugar regulation effects are also SMF direction dependent. Although the Lorenz forces exerted on negatively charged moving DNA can explain some of the phenotypes, we believe there must be other unraveled mechanisms underlying the SMF direction dependent bioeffects.

In summary, our study revealed that a moderate SMF with a downward direction could alleviate ALD through reduced ROS, oxidative stress, and inflammation. Unexpectedly, the upward SMF setting in our study may be a taboo for heavy drinkers, which might be a potential precaution to some open MRI operations, which usually have upright magnetic field of ∼0.5 T. Whether other magnetic settings, for example, different magnetic field flux density and distribution, will provide improved beneficial effects or more detrimental effects, still need further investigations. However, we found that a relatively simple magnetic setting could bring obvious benefits for both light and heavy drinking mice by reducing the oxidative stress. Our study illustrates the potential to develop magnetic field as a physical tool for preventive and/or clinical therapeutic treatment strategy of ALD, as well as other oxidative stress-related health applications in the future.

## Methods

### Mice model

Eight-week-old male BALB/c mice purchased from Nanjing biomedical Research Institute of Nanjing University (Nanjing, China) were used in this study. All mice were randomly divided into EtOH-fed groups (sham, upward and downward SMF; ethanol diet) and pair-fed groups (sham, upward and downward SMFs; maltodextrin diet). Lieber-DeCarli diets (product L10015 and product L10016) were prepared fresh daily and fed ad libitum to mice using a short-term (chronic EtOH-fed mice model with light drinking) and long-term (acute EtOH-fed mice model with heavy drinking) ethanol feeding protocol of ALD according to the previous report (Wang et al., 2020). The formulas of diets were formulated by Research Diets. The mice were exposed to sham, upward and downward SMFs for 3 weeks (chronic model) or 6 weeks (acute model). Acute EtOH-fed mice model: the diet containing ethanol were designed as 1% ethanol for 1 week, 3% ethanol for 1 week, 5% ethanol for 1 week, 6% ethanol for 3 weeks. Chronic EtOH-fed mice model: the diet containing ethanol were designed as 1%, 2% and 3% ethanol for 1 week, respectively. The diet consumption and body weight were recorded during the whole experiment. After the mice were executed, serum and organs were collected and stored at -80°C.

All animal experiments were conducted according to the Guide for the Care and Use of Laboratory Animals of the National Institutes of Health and carried out strictly in accordance with the approved institutional animal care and use committee (IACUC) of Hefei Institutes of Physical Science, Chinese Academy of Science (Hefei, China). The protocol was approved by the Committee on the Ethics of Animal Experiments of Hefei Institutes of Physical Science, Chinese Academy of Science (Permit Number: SW YX-DW-2021-54).

### Magnetic field setup

The upward and downward magnetic fields were provided by permanent magnet plates (length × width × height = 250 mm ×160 mm × 45 mm). Each magnetic plate was composed of 12 Neodymium magnet cubes (length × width × height = 60 mm × 50 mm × 30 mm) that were purchased from Sans (Nanjing, China). These magnet plates can produce a relatively uniform static magnetic field at each horizontal plane. Experimental groups mice were continuously espoused to SMFs for 12 h per day during the light and dark cycle. To measure the distributions of magnetic field at the mice location, a magnet analyzer (FE-2100RD, Forever Elegance, China) was used to scan the planes at 2 cm above the magnets. The average magnetic flux density of the magnets at this horizontal plane is approximately 0.1 T. Additionally, cultured cells were also exposed to Neodymium magnets (N38, length × width × height = 60 mm × 50 mm × 30 mm) with the surface magnetic flux density of 0.4 T.

### Physiological index detection

The physiological condition of mice was monitored by the small animal vital signs monitor (STARR Life Science, USA). The signal sensor was placed on the necks of the mice, their hair can affect signal reception. Therefore, BALB/c mice with white hair were selected to perform the experiment. Mice were placed in the cage and monitored for 6 min. The breath rate, pulse distention, heart rate and arterial O_2_ were calculated using Mouse-ox plus software (STARR Life Science, USA).

### Hematoxylin–eosin (H&E) staining

At the end of the experiment, all mice were dissected to collect organs including heart, liver, spleen, lung, kidney. Then organs were fixed and processed with formalin to obtain 5-μm-thick sections and stained with H&E. Five random areas were examined in each section.

### TUNEL assay

Terminal deoxynucleotidyl transferase dUTP nick-end labeling (TUNEL) detection kit (C1098, Beyotime, China) was used to detect apoptotic cells in liver tissue. 3,3’-diaminobenzidine (DAB) reagent stained and incubated at room temperature for 5-30 min. Liver sections of TUNEL staining were quantitatively analyzed by counting TUNEL positive cells from at least 5 microscopic areas.

### Oil red O staining

Oil red O staining solution (R23104, Baso, China) was used to detect lipid accumulation in liver sections and human hepatocyte HL7702 cells. HL7702 cells were seeded in 35 mm cultured dishes, treated with ethanol (0 mM, 85 mM, 170 mM, 340 mM) or magnetic fields (sham, upward or downward SMFs) for 24 h. Oil red O mixed with ddH_2_O at the ratio of 3:2 and was left at room temperature for 10 min. Oil red O staining solution and 70% propylene glycol were added with agitation for 5-10 min, followed by washing in running water for 30 s. Hematoxylin solution was used for staining for 2 min, and then washed off with running water for 5 min. At last, stained sections and cells were examined by microscopy.

### Blood chemistry and tissue MDA tests

The blood samples were collected in 1.5 mL centrifuge tube at 4°C, 3500×g for 15 min to collect serum, which were analyzed by an automated biochemical analyzer (HITACHI 7020, Japan). Lipid peroxidation MDA assay kit (S0131S, Beyotime, China) were used to detect the MDA concentration in liver tissues according to the manufacturer’s instructions.

### Immunohistochemistry

Liver section immunohistochemistry was performed using the antibodies for NRF2 (abs130481, ABSIN), MPO (79623S, CST), Ki67 (27309-1-AP, Proteintech), Top2A (202331-1-AP, Proteintech) and F4/80 (70076S, CST). All the steps were performed according to the manufacturer’s instructions. Quantitation of staining was evaluated by ImageJ analysis of at least 5 microscopic areas.

### Cell number counting and apoptosis analysis by Flow cytometry

Human hepatocyte HL7702 cells (ATCC) were exposed to alcohol and magnetic fields before they were collected in 1.5 mL centrifuge tube by 0.25% trypsin digestion. Cell number were counted by flow cytometry. Additionally, cultured cells were exposed to alcohol and SMFs for 24 h. Annexin V-FITC apoptosis detection kit (BD Biosciences, 0076884, USA) was used to analyze hepatocyte apoptosis according to manufacturer’s protocol. Cell apoptosis was detected by flow cytometry.

### Western blot analysis

The protein isolations were collected with protease inhibitors from liver tissues and cultured cells, and western blotting were performed as previously described. The proteins transferred to nitrocellulose membranes were incubated with primary antibodies NRF2 (12721S, CST), BCL2 (26295S, CST), cleaved-caspase3 (9664S, CST), caspase3 (9662S, CST), anti-tubulin (2128S, CST) and anti-GAPDH (5174S, CST). ImageJ software was used to quantitatively analyze the expression of proteins.

### RT-PCR and QPCR

Mice exposed to the short-term alcoholic diet were executed to collect liver tissues. And then liver tissues were polished on ice and were used to extract RNA by a RNAeasy™ animal RNA isolation kit with spin column (R0027, Beyotime). Novoscript^R^ plus all-in-one 1^st^ strand cDNA synthesis supermix (E047-01A, Novoprotein) was used to RNA reverse transcription, and Novostart^R^SYBR qPCR supermix plus (E096-01A, Novoprotein) were used to amplify target gene under the action of specific primers. The detailed primers sequences were shown in table S1.

### Cellular Reactive Oxygen Species (ROS) detection

HL7702 cells (4 ×10^4^ cells/mL) were seeded in 35 mm culture dish and supplied with complete medium. After attachment, the medium was supplemented with different concentrations of ethanol (0 mM, 85 mM, 170 mM, 340 mM) and/or exposed to sham, upward or downward SMFs for 24 h. ROS detection kit (D6883, Sigma) containing 2’,7’-dichlorodihydrofluorescein diacetate (DCFH-DA) was used to detect cellular ROS. DCFH-DA can freely penetrate the cell membrane and is hydrolyzed by esterase in the cell to produce DCFH that cannot pass through the cell membrane, while ROS can oxidize non-fluorescent DCFH to produce fluorescent DCF. Cultured cells were incubated with 10 µM DCFH-DA at 37°C for 30 min. Green fluorescence can be observed by a fluorescence microscope, and the intensity of green fluorescence is directly proportional to the level of cellular ROS. Meanwhile, flow cytometry and confocal fluorescence microscope were used to evaluate the intensity of fluorescent DCF.

### EPR (Electron Paramagnetic Resonance) experiment

500 μL 0.2 M H_2_O_2_ solution was made by mixing 10 μL H_2_O_2_ (10 M, AR≥30%) and 490 μL PBS buffer (137 mM NaCl, 2.7 mM KCl, 10 mM Na_2_HPO_4_, 1.8 mM KH_2_PO_4_), and was then exposed to the SMFs for 24 h. 5,5-dimethyl-1-pyrroline N-oxide (DMPO) (92688, Sigma) was added to the solution 5 min before EPR experiments. The free radical levels from equal amount solutions from each condition were tested by the EPR device (Bruker EMX plus 10/12, Switzerland) equipped with Oxford ESR910 Liquid Helium cryostat.

### DNA synthesis assay

HL7702 cells (4 ×10^4^ cells/mL) were seeded in 35 mm dish and synchronized with 2.5 mM thymidine for 16 h, and then treated with 10 μM BrdU for 8 h when they were exposed to SMF or sham at indicated time point. Then, all cells were fixed by 70 % ethanol for 12 h and washed by PBS, resuspended by 2 M HCl and incubated on the rotator for 30 min at room temperature, centrifuged and resuspended by 0.1 M Na_2_B_4_O_7_ (Ph 8.5) at room temperature for 10 min before they were washed by PBS. Finally, the cells were incubated with anti-BrdU antibody for 2.5 h and the secondary Alexa Fluor 488 conjugated antibody for 1.5 h. All cells were analyzed by flow cytometry.

### Statistical analysis

All statistical analysis was performed using GraphPad Prism version 8. Date from the experiments were showed as the means ± SEM. The *P*-values were calculated using the one-way or two-way analysis of variance (ANOVA) with Bonferroni correction or two-tailed unpaired *t*-test for comparisons. *P* < 0.05 was considered statistically significant.

## Acknowledgements

We thank Shu-Tong Maggie Wang for cartoon illustration. A portion of this work was performed on the Steady High Magnetic Field Facilities, High Magnetic Field Laboratory, CAS.

## Funding

The work was supported by the National Key Research and Development Program of China (Xin Zhang), National Natural Science Foundation of China 52007185 (Xiaofei Tian), HFIPS Director’s Fund YZJJ2020QN26 (Xinmiao Ji) and Heye Health Technology Foundation HYJJ20190801 (Xin Zhang).

## Author contributions

Xin Zhang and Chao Song designed the study, analyzed the data, and wrote the manuscript with input from the other authors. Chao Song, Hanxiao Chen performed experiments and analyzed data. Biao Yu, Lei Zhang, Chuanlin Feng, Xingxing Yang, Xiaofei Tian, Junjun Wang, Yixiang Fang and Xinmiao Ji contributed to editing, revising, and finalizing the manuscript. Hua Wang and Can Xie contributed with discussion, data abstraction, and data interpretation. All authors have read and agreed to the published version of the manuscript.

## Competing interests

The authors declare no competing interests.

## Data availability

All data needed to evaluate the conclusions in the paper are present in the paper and/or the Supplementary Materials. Additional data related to this paper may be requested from the authors.

## Source data

Figure 1-source data 1: Dataset values for Figure 1D.

Figure 1-source data 2: Dataset values for Figure 1F.

Figure 1-source data 3: Raw western blot images for Figure 1G.

Figure 1-figure supplement 1-source data 1: Dataset values for Figure 1-figure supplement 1A.

Figure 1-figure supplement 1-source data 2: Dataset values for Figure 1-figure supplement 1B.

Figure 2-source data 1: Dataset values for Figure 2B.

Figure 2-source data 2: Dataset values for Figure 2C.

Figure 3-source data 1: Dataset values for Figure 3B.

Figure 3-source data 2: Dataset values for Figure 3D.

Figure 3-source data 3: Dataset values for Figure 3E.

Figure 3-source data 4: Dataset values for Figure 3G.

Figure 3-source data 5: Dataset values for Figure 3H.

Figure 4-source data 1: Dataset values for Figure 4C.

Figure 4-source data 2: Dataset values for Figure 4D.

Figure 4-source data 3: Dataset values for Figure 4E.

Figure 4-source data 4: Dataset values for Figure 4F.

Figure 4-source data 5: Raw western blot images for Figure 4G.

Figure 4-source data 6: Dataset values for Figure 4H.

Figure 4-source data 7: Dataset values for Figure 4I.

Figure 5-source data 1: Dataset values for Figure 5B.

Figure 5-source data 2: Dataset values for Figure 5D.

Figure 5-source data 3: Dataset values for Figure 5E.

Figure 6-source data 1: Dataset values for Figure 6C and 6E.

Figure 7-source data 1: Dataset values for Figure 7C.

Figure 7-source data 2: Dataset values for Figure 7D.

Figure 7-source data 3: Dataset values for Figure 7H.

Figure 8-figure supplement 7-source data 1: Dataset values for Figure 8-figure supplement 7.

## Notes

### Competing Interest Statement

The authors have declared no competing interest.

## References

Alarcón-Sánchez BR, Guerrero-Escalera D, Rosas-Madrigal S, Ivette Aparicio-Bautista D, Reyes-Gordillo K, Lakshman MR, Ortiz-Fernández A, Quezada H, Medina-Contreras Ó, Villa-Treviño S, Isael Pérez-Carreón J, Arellanes-Robledo J. Nucleoredoxin interaction with flightless-I/actin complex is differentially altered in alcoholic liver disease. Basic Clin Pharmacol Toxicol. 2020;127: 389–404. doi: 10.1111/bcpt.13451.

Albano E. Alcohol, oxidative stress and free radical damage. Proc Nutr Soc. 2006;65:278–290. doi: 10.1079/pns2006496.

Barnes F, Greenebaum B. Role of radical pairs and feedback in weak radio frequency field effects on biological systems. Environ Res. 2018;163:165–170. doi:10.1016/j.envres.2018.01.038.

Bhandari R, Khaliq K, Ravat V, Kaur P, Patel RS. Chronic Alcoholic Liver Disease and Mortality Risk in Spontaneous Bacterial Peritonitis: Analysis of 6,530 Hospitalizations. Cureus. 2020;12:e8189. doi: 10.7759/cureus.8189.

Carter CS, Huang SC, Searby CC, Cassaidy B, Miller MJ, Grzesik WJ, Piorczynski TB, Pak TK, Walsh SA, Acevedo M, Zhang Q, Mapuskar KA, Milne GL, Hinton AO Jr, Guo DF, Weiss R, Bradberry K, Taylor EB, Rauckhorst AJ, Dick DW, Akurathi V, Falls-Hubert KC, Wagner BA, Carter WA, Wang K, Norris AW, Rahmouni K, Buettner GR, Hansen JM, Spitz DR, Abel ED, Sheffield VC. Exposure to Static Magnetic and Electric Fields Treats Type 2 Diabetes. Cell Metab. 2020;32:561-574.e7. doi:10.1016/j.cmet.2020.09.012.

Cederbaum AI. Alcohol metabolism. Clin Liver Dis. 2012;16:667–685. doi:10.1016/j.cld.2012.08.002.

Das SK, Vasudevan DM. Alcohol-induced oxidative stress. Life Sci. 2007;81:177–187. doi:10.1016//j.lfs.2007.05.005.

de Labry LO, Glynn RJ, Levenson MR, Hermos JA, LoCastro JS, Vokonas PS. Alcohol consumption and mortality in an American male population: recovering the U-shaped curve--findings from the normative Aging Study. J Stud Alcohol. 1992;53:25–32. doi: 10.15288/jsa.1992.53.25.

Esser MB, Sherk A, Liu Y, Naimi TS, Stockwell T, Stahre M, Kanny D, Landen M, Saitz R, Brewer RD. Deaths and Years of Potential Life Lost From Excessive Alcohol Use-United States, 2011-2015. Morb Mortal Wkly Rep. 2020;69:981–987. doi: 10.15585/mmwr.mm6930a1.

Grissom CB. Magnetic-Field Effects in Biology - a Survey of Possible Mechanisms with Emphasis on Radical-Pair Recombination. Chemical Reviews. 1995;95:3–24. doi: 10.1021/cr00033a001.

Gurhan H, Bruzon R, Kandala S, Greenebaum B, Barnes F. Effects Induced by a Weak Static Magnetic Field of Different Intensities on HT-1080 Fibrosarcoma Cells. Bioelectromagnetics. 2021;42:212–223. doi: 10.1002/bem.22332.

Ikeya N, Woodward JR. Cellular autofluorescence is magnetic field sensitive. Proc Natl Acad Sci U S A. 2021;118:e2018043118. doi: 10.1073/pnas.2018043118.

Liang R, Liu A, Perumpail RB, Wong RJ, Ahmed A. Advances in alcoholic liver disease: An update on alcoholic hepatitis. World J Gastroenterol. 2015;21:11893–903. doi: 10.3748/wjg.v21.i42.11893.

Lieber CS, DeCarli LM, Sorrell MF. Experimental methods of ethanol administration. Hepatology. 1989;10:501–510. doi: 10.1002/hep.1840100417. PMID: 2673971.

Mansouri A, Gattolliat CH, Asselah T. Mitochondrial Dysfunction and Signaling in Chronic Liver Diseases. Gastroenterology. 2018;155:629–647. doi: 10.1053/j.gastro.2018.06.083.

Marmot MG, Rose G, Shipley MJ, Thomas BJ. Alcohol and mortality: a U-shaped curve. The Lancet. 1981;1:580–573. doi: 10.1016/s0140-6736(81)92032-8.

Osna NA, Kharbanda KK. Multi-Organ Alcohol-Related Damage: Mechanisms and Treatment. Biomolecules. 2016;6:20. doi: 10.3390/biom6020020.

Parker R, Aithal GP, Becker U, Gleeson D, Masson S, Wyatt JI, Rowe IA; WALDO study group. Natural history of histologically proven alcohol-related liver disease: A systematic review. J Hepatol. 2019;71:586–593. doi: 10.1016/j.jhep.2019.05.020.

Rehm J, Samokhvalov AV, Shield KD. Global burden of alcoholic liver diseases. J Hepatol. 2013;59:1 60–168. doi: 10.1016/j.jhep.2013.03.007.

Seitz HK, Bataller R, Cortez-Pinto H, Gao B, Gual A, Lackner C, Mathurin P, Mueller S, Szabo G, Tsukamoto H. Alcoholic liver disease. Nat Rev Dis Primers. 2018;4:16. doi: 10.1038/s41572-018-0014-7.

Sha’fie MSA, Rathakrishnan S, Hazanol IN, Dali MHI, Khayat ME, Ahmad S, Hussin Y, Alitheen NB, Jiang LH, Syed Mortadza SA. Ethanol Induces Microglial Cell Death via the NOX/ROS/PARP/TRPM2 Signalling Pathway. Antioxidants (Basel). 2020;9:12 53. doi: 10.3390/antiox9121253.

Shaper AG, Wannamethee G, Walker M. Alcohol and mortality in British men: explaining the U-shaped curve. The Lancet. 1988;2:1267–1273. doi: 10.10.16/s0140-6736(88)92890-5.

Singal AK, Bataller R, Ahn J, Kamath PS, Shah VH. ACG Clinical Guideline: Alcoholic Liver Disease. Am J Gastroenterol. 2018;113:175–194. doi: 10.1038/ajg.2017.469.

Sy-Janairo ML Y, Cua IH. Association of metabolic-associated fatty liver disease and risk of severe coronavirus disease 2019 illness. JGH Open. 2020;5:4–10. doi: 10.1002/jgh3.12465.

Thursz M, Kamath PS, Mathurin P, Szabo G, Shah VH. Alcohol-related liver disease: Areas of consensus, unmet needs and opportunities for further study. J Hepatol. 2019;70:521–530. doi:10.1016/j.jhep.2018.10.041.

Timmel CR, Till U, Brocklehurst B, McLauchlan KA, Hore PJ. The effects of weak magnetic fields on radical recombination reactions. Molecular Physics. 1998;95:71–89. doi: 10.1080/00268979809483134.

Van Huizen AV, Morton JM, Kinsey LJ, Von Kannon DG, Saad MA, Birkholz TR, Czajka JM, Cyrus J, Barnes FS, Beane WS. Weak magnetic fields alter stem cell-mediated growth. Sci Adv. 2019;5:eaau7201. doi: 10.1126/sciadv.aau7201.

Wang H, Zhou H, Zhang Q, Poulsen KL, Taylor V, McMullen MR, Czarnecki D, Dasarathy D, Yu M, Liao Y, Allende DS, Chen X, Hong L, Zhao J, Yang J, Nagy LE, Li X. Inhibition of IRAK4 kinase activity improves ethanol-induced liver injury in mice. J Hepatol. 2020;73:1470–1481. doi: 10.1016/j.jhep.2020.07.016.

Wang H, Zhang X. Magnetic Fields and Reactive Oxygen Species. Int J Mol Sci. 2017;18:2175. doi: 10.3390/ijms18102175.

Wang H, Zhang X. ROS Reduction Does Not Decrease the Anticancer Efficacy of X-Ray in Two Breast Cancer Cell Lines. Oxid Med Cell Longev. 2019;2019:3782074. doi: 10.1155/2019/3782074.

Wang Z, Dou X, Li S, Zhang X, Sun X, Zhou Z, Song Z. Nuclear factor (erythroid-derived 2)-like 2 activation-induced hepatic very-low-density lipoprotein receptor overexpression in response to oxidative stress contributes to alcoholic liver disease in mice. Hepatology. 2014;59:1381–1392. doi: 10.1002/hep.26912.

Williams R, Alessi C, Alexander G, Allison M, Aspinall R, Batterham RL, Bhala N, Day N, Dhawan A, Drummond C, Ferguson J, Foster G, Gilmore I, Goldacre R, Gordon H, Henn C, Kelly D, MacGilchrist A, McCorry R, McDougall N, Mirza Z, Moriarty K, Newsome P, Pinder R, Roberts S, Rutter H, Ryder S, Samyn M, Severi K, Sheron N, Thorburn D, Verne J, Williams J, Yeoman A. New dimensions for hospital services and early detection of disease: a Review from the Lancet Commission into liver disease in the UK. The Lancet. 2021;397:1770–1780. doi: 10.1016/S0140-6736(20)32396-5.

Witkiewitz K, Litten RZ, Leggio L. Advances in the science and treatment of alcohol use disorder. Sci Adv. 2019;5:eaax4043. doi: 10.1126/sciadv.aax4043. PMID: 3157982.

Wu D, Cederbaum AI. Alcohol, oxidative stress, and free radical damage. Alcohol Res Health. 2003;27:277–284.

Xiao J, Wang F, Wong NK, Lv Y, Liu Y, Zhong J, Chen S, Li W, Koike K, Liu X, Wang H. Epidemiological Realities of Alcoholic Liver Disease: Global Burden, Research Trends, and Therapeutic Promise. Gene Expr. 2020;20:105–118. doi: 10.3727/105221620X15952664091823.

Yang X, Li Z, Polyakova T, Dejneka A, Zablotskii V, Zhang X. Effect of static magnetic field on DNA synthesis: The interplay between DNA chirality and magnetic field left-right asymmetry. FASEB Bioadv. 2020;2:254–263. doi: 10.1096/fba.2019-00045.

Yang X, Song C, Zhang L, Wang J, Yu X, Yu B, Zablotskii V, Zhang X. An upward 9.4 T static magnetic field inhibits DNA synthesis and increases ROS-P53 to suppress lung cancer growth. Transl Oncol. 2021;14:101103. doi: 10.1016/j.tranon.2021.101103.

Yu B, Choi B, Li W, Kim DH. Magnetic field boosted ferroptosis-like cell death and responsive MRI using hybrid vesicles for cancer immunotherapy. Nat Commun. 2020;11:3637. doi: 10.1038/s41467-020-17380-5.

Yu B, Liu J, Cheng J, Zhang L, Song C, Fang YX, Lu Y, Zhang X. A Static Magnetic Field Improves Iron Metabolism and Prevents High-Fat-Diet/Streptozocin-Induced Diabetes. The Innovation. 2021;2:100077. doi:10.1016-j.xinn.2021.100077.

